# Mapping Trends in Insecticide Resistance Phenotypes in African Malaria Vectors

**DOI:** 10.1101/2020.01.06.895656

**Authors:** PA Hancock, CJM Hendriks, J-A Tangena, H Gibson, J Hemingway, M Coleman, PW Gething, E Cameron, S Bhatt, CL Moyes

## Abstract

Mitigating the threat of insecticide resistance in African malaria vector populations requires comprehensive information about where resistance occurs, to what degree, and how this has changed over time. Estimating these trends is complicated by the sparse, heterogeneous distribution of observations of resistance phenotypes in field populations. We use 6423 observations of the prevalence of resistance to the most important vector control insecticides to inform a Bayesian geostatistical ensemble modelling approach, generating fine-scale predictive maps of resistance phenotypes in mosquitoes from the *Anopheles gambiae* complex across Africa. Our models are informed by a suite of 111 predictor variables describing potential drivers of selection for resistance. Our maps show alarming increases in the prevalence of resistance to pyrethroids and DDT across Sub-Saharan Africa from 2005-2017 as well as substantial spatial variation in resistance trends.

## INTRODUCTION

Insecticide resistance in African malaria vector populations has serious consequences for malaria prevention. Long-lasting insecticide-treated bednets (LLINs) have achieved substantial reductions in malaria prevalence thus far in Africa^1^, but the number of insecticides currently available for use in LLINs is very limited. Until recently, pyrethroids were the only class approved for use in LLINs and recently launched new generation nets still use pyrethroids in combination with either an insect growth regulator, a pyrrole, or a synergist that inhibits the primary metabolic mechanism of pyrethroid resistance within mosquitoes^2, 3^. A wider range of options is available for indoor residual spraying (IRS), but pyrethroids are less expensive than many alternatives and are still used for IRS in malaria-endemic Sub-Saharan African countries^4, 5^.

Although there is evidence that pyrethroid resistance in African malaria vector populations is increasing^6, 7^, the wide array of field studies that are available do not provide a spatially-comprehensive time series of resistance trends^8^. Quantifying these trends will improve our understanding of the historical spread of resistance and assist in designing insecticide resistance management strategies^9^. Comprehensive spatiotemporal analyses of resistance are also necessary to facilitate its inclusion in epidemiological models of malaria that inform decision-making at national and global levels^9^. Efforts to estimate trends in insecticide resistance are impeded by limitations associated with the available observations of resistance phenotypes in field mosquito populations. Observations from standardized susceptibility tests, which indicate the prevalence of phenotypic resistance in field populations, cover a wide geographic area and span several decades^8, 10^. However, the spatial coverage of this data is sparse and heterogeneous, and resistance has rarely been monitored consistently over time, meaning that very few time series are available^9^. Moreover, these susceptibility tests have a large measurement error, and replication is required to robustly estimate resistance phenotypes.

Our capacity to understand and predict insecticide resistance can benefit from considering the variables that may influence selection for resistance. Sources of insecticides in the environment include the application of insecticide-based vector control interventions for public health, such as LLINs and IRS, and the application of agricultural insecticides, which include the same insecticide classes as those used in vector control^11^. Several studies have demonstrated a local increase in insecticide resistance in field mosquito populations following the implementation of LLINs, IRS, or both^12, 13, 14, 15, 16^ although in other locations evidence of higher resistance after the introduction these interventions was not found^12, 17^. Associations between agricultural pesticide use and insecticide resistance have also been found^11, 18^, and there is evidence that pesticide contamination of water bodies is a source of selection pressure for resistance acting on mosquito larvae^19^. Relationships between resistance and drivers of selection will, however, vary geographically depending on population structure^20, 21^. Genetic mechanisms of resistance also differ across mosquito species^15, 20^, and even closely-related mosquito species have different ecological niches^22, 23^, as well as different blood feeding behaviour and preferences, meaning that they are likely to experience differences in insecticide exposure^24^.

To develop predictive models of insecticide resistance in field populations that can represent variable, nonlinear interactions with environmental, biological and genetic variables, we utilise an ensemble modelling approach. The approach exploits the multi-faceted strengths of different modelling methodologies, using machine-learning methods to extract predictive power from a set of covariates, and then allowing a Bayesian geostatistical Gaussian process to model the autocorrelated residual variation^25^. Bayesian geostatistical models provide a robust model of residual autocorrelation that can be applied to spatiotemporal data with a heterogeneous sampling distribution^26^. Their application to observations from insecticide susceptibility tests conducted over a range of locations across Africa has previously demonstrated broad-scale associations between resistance to different types pyrethroids, as well as the organochlorine DDT^27^. The models developed in this study exploit these associations in resistance across different insecticides to improve resistance predictions for individual insecticide types.

Using a database containing the results of standard insecticide susceptibility tests performed on mosquito samples collected throughout Africa^8^, we extracted the results of 6423 tests conducted on samples from the *Anopheles gambiae* species complex, which are among the most important African malaria vectors. We used this data set in our model ensemble to quantify variation in the prevalence of resistance to pyrethroids and DDT over the period 2005-2017 by developing a series of predictive maps. Our models are informed by a suite of potential explanatory variables describing the coverage of insecticide-based vector control interventions, agriculture and other types of land cover, climate, processes determining the environmental fate of pesticides, and the distribution of the sibling species that make up the *An. gambiae* complex. Our results show dramatic changes in insecticide resistance phenotypes in malaria vector populations across Africa over a thirteen-year period, and identify variables that were important in shaping these predictions.

## RESULTS

### Spatiotemporal trends in the prevalence of insecticide resistance

#### Pyrethroid resistance

We investigated spatiotemporal trends in the prevalence of phenotypic resistance in the *An. gambiae* complex to four pyrethroids: deltamethrin, permethrin, lambda-cyhalothrin and alpha-cypermethrin. Due to the lack of observations from central Africa, we partitioned the data into two separate spatial regions covering western and eastern parts of the continent, and analysed each data subset independently by fitting separate models (see Methods). In west Africa, predicted mean prevalence of resistance to all pyrethroids increased dramatically over the period 2005-2017 (Figs. 1, 2 and Supplementary Figs. 1, 2 & 3). Predicted mean proportional mortality to deltamethrin was below 0.9 (the WHO threshold for confirmed resistance) across 15% (95% credible interval (CI) = 13-17%) of the west region in 2005, and across 98% (CI=96.6-98.7%) of the region in 2017 (Fig. 2 and see Supplementary Fig. 8 for the trends for individual countries). These changes in resistance were spatially heterogeneous (Fig. 1). Increases in resistance to deltamethrin over the period, in terms of the reductions in the predicted mean proportional mortality, were greatest in northern Liberia (Fig. 1D, line A), central Cote d’Ivoire (Fig. 1D, line B), the area surrounding the border between Burkina Faso, Cote d’Ivoire and Ghana (Fig. 1D, line C), southern Ghana (Fig. 1D, line D), and northern Gabon (Fig. 1D, line E). In these regions, resistance to deltamethrin in 2017 was particularly high (with a mean proportional mortality below 0.3 (CI <0.4).

**Figure 1.**
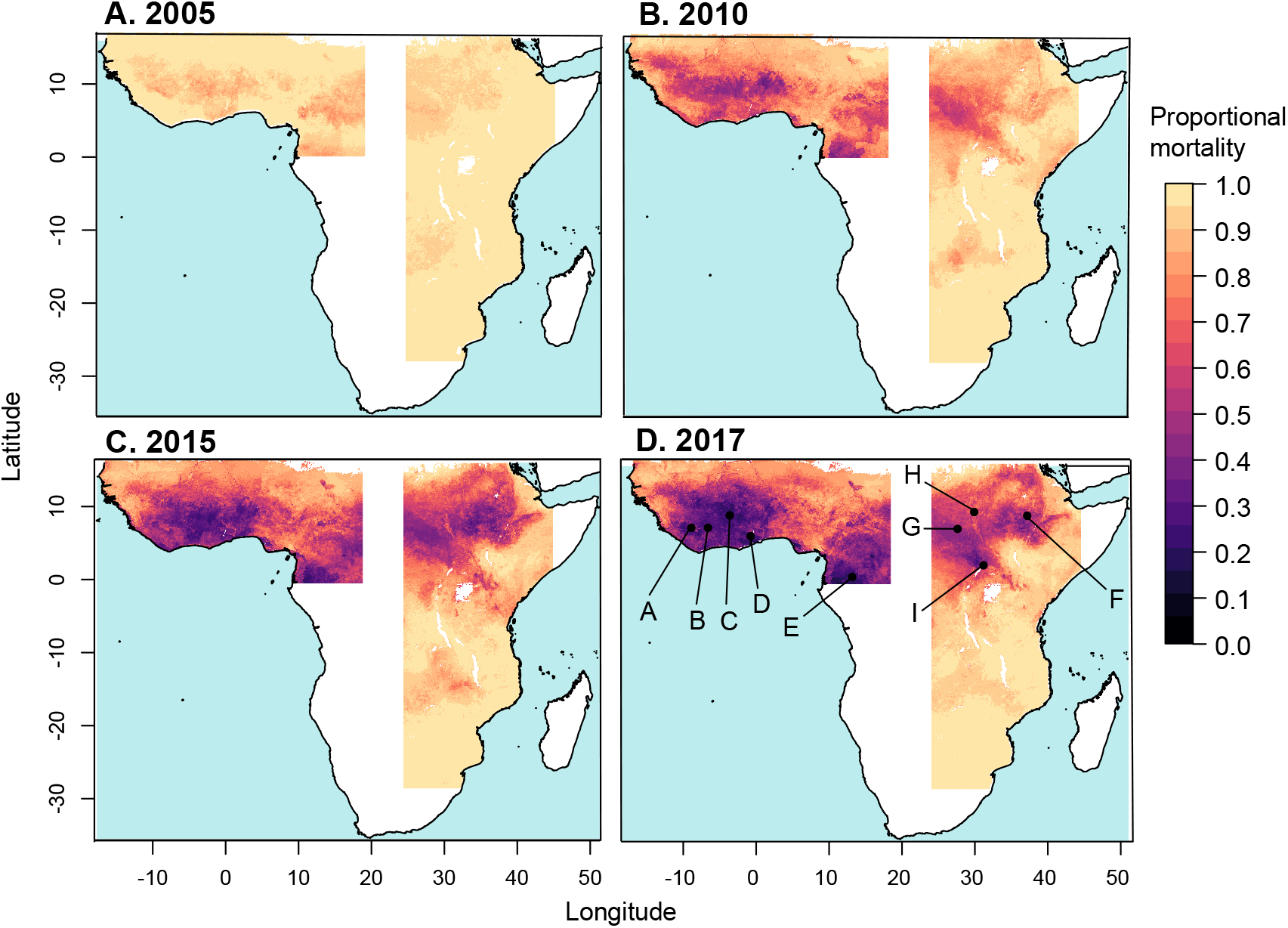
Predicted mean proportional mortality to deltamethrin across the west and east regions. A. 2005; B. 2010; C. 2015; D. 2017.

In east Africa, the prevalence of pyrethroid resistance also increased over the period 2005-2017, albeit at a lesser rate than that in the west region (Figs. 1 & 2). Predicted mean proportional mortality to deltamethrin was below 0.9 across 9% (CI=3-17%) of the east region in 2005 and across 45% (CI=38-51%) of the region in 2017 (Fig. 2 and see Supplementary Fig. 8 for the trends for individual countries). The greatest increases in pyrethroid resistance over the period occurred in the northern part of the region, in the area from central Ethopia (Fig. 1D, line F) westward across most of South Sudan (Fig. 1D, line G), and extending into southern Sudan (Fig. 1D, line H) and northern Uganda (Fig. 1D, line I). Across most of this area, mean mortality to deltamethrin in 2017 was below 0.5 (CI < 0.75). Resistance to deltamethrin increased to a lesser extent in central and southern Uganda, western Kenya, eastern Ethopia and coastal Tanzania, with predicted mean mortalities of between 0.6-0.8 in these areas in 2017. In areas further south, differences in predicted resistance over the time period were relatively slight, with mean mortalities changing by less than 0.15 from 2005-2017 within Malawi, Mozambique, Zimbabwe, and those parts of Zambia, Botswana and South Africa that were included in the model. Similar spatiotemporal trends across the west and east regions occurred in predicted mean resistance to permethrin, lambda-cyhalothrin and alpha-cypermethrin (Supplementary Figs. 1, 2 & 3).

**Figure 2.**
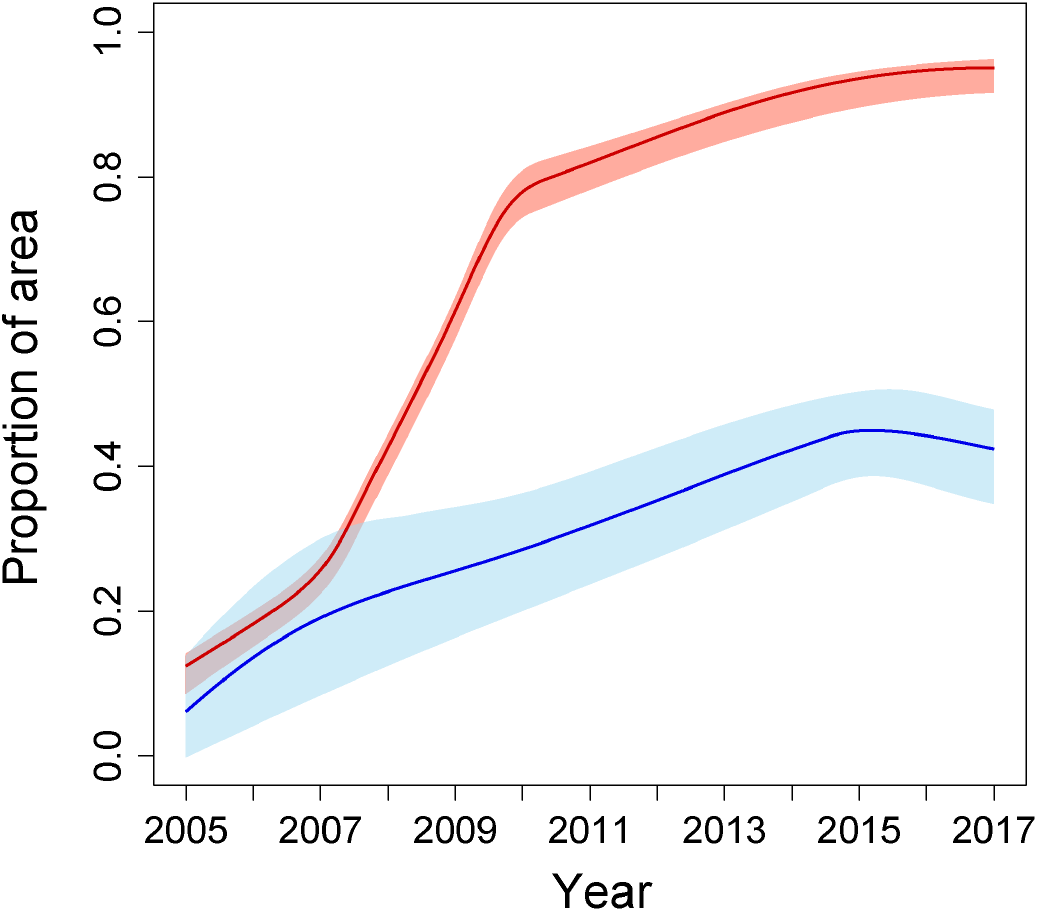
The proportion of the area with a predicted mean mortality to deltamethrin of less than 0.9, for the west region (red line) and the east region (blue line). Red and blue shaded areas indicate the 95% credible interval of the predicted proportion of pixels for the west and east regions, respectively.

#### DDT resistance

Predicted mean resistance to DDT at the start of the period (in 2005) was more widespread in comparison to pyrethroid resistance, and also increased throughout the region from 2005-2017 (Fig. 3 & 4). In the west region, predicted mean proportional mortality to DDT was below 0.9 across 53% (CI = 47-59%) of the west region in 2005, and across 97% (CI=92.7-99%) of the region in 2017 (Fig. 4). Increases in resistance to DDT over the period were greatest in the area surrounding the border between Liberia and Guinea (Fig. 3D, line A), southern Mali (Fig. 3D, line B), and central Burkina Faso (Fig. 3D, line C). The east region showed a weaker increase in predicted mean resistance to DDT over the period 2005-2017 in comparison to that occurring in the west region. Predicted mean proportional mortality was below 0.9 across 32% (CI=21-44%) of the east region in 2005, and across 45% (CI=39-51%) in 2017. Increases in DDT resistance over the period were greatest in central South Sudan (Fig. 3D, line D).

**Figure 3.**
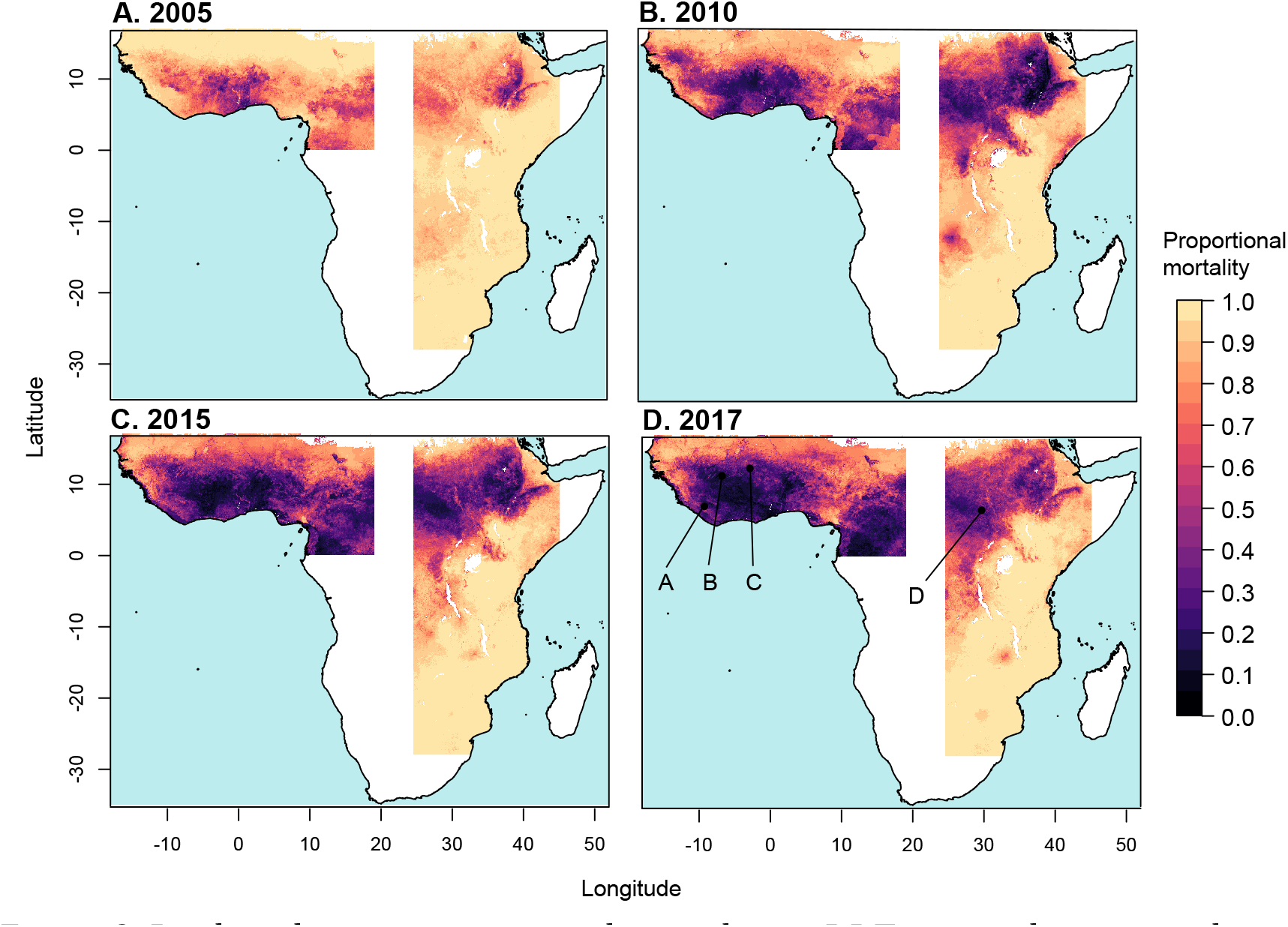
Predicted mean proportional mortality to DDT across the west and east regions. A. 2005; B. 2010; C. 2015; D. 2017.

**Figure 4.**
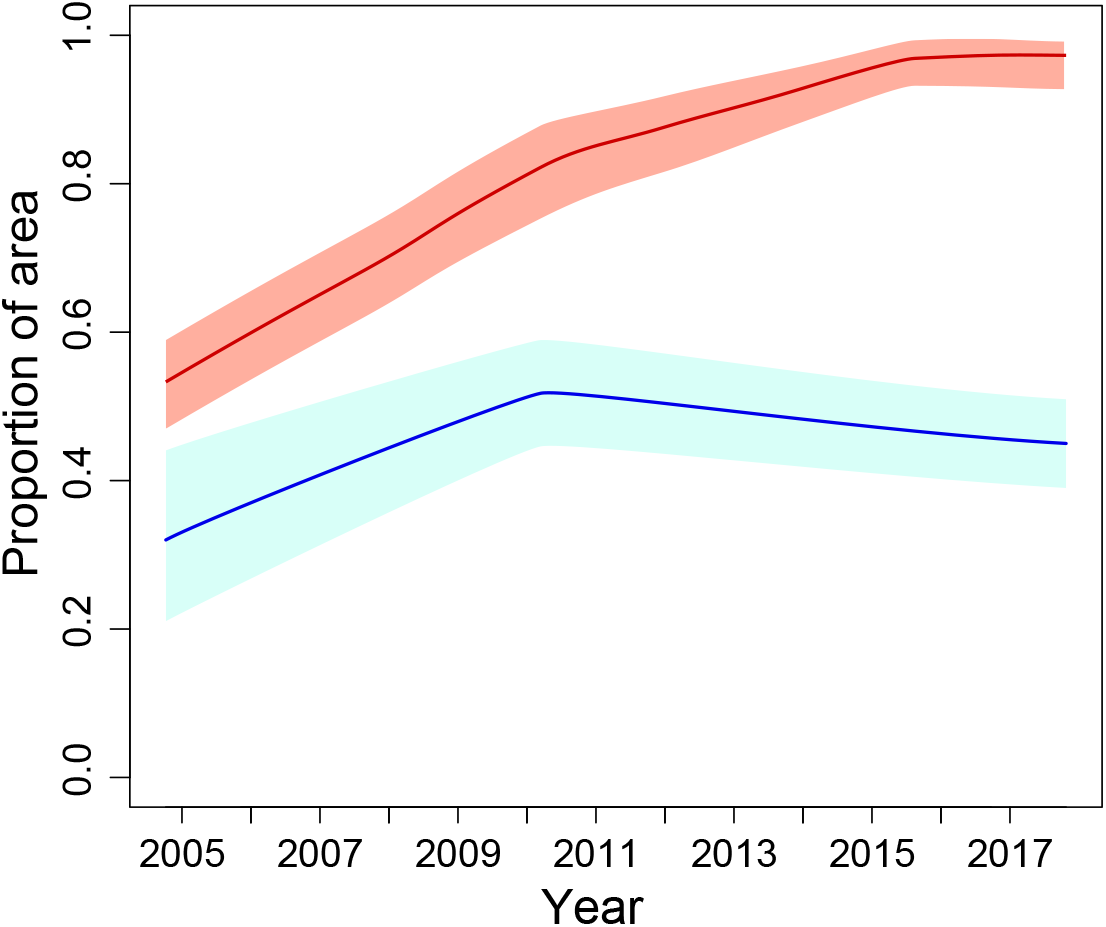
The proportion of the area with a predicted mean mortality to DDT of less than 0.9, for the west region (red line) and the east region (blue line). Red and blue shaded areas indicate the 95% credible interval of the predicted proportion of pixels for the west and east regions, respectively.

### Assessing prediction accuracy

We performed 10-fold out-of-sample validation on the model ensemble to assess the accuracy of predicted mean prevalence of resistance. Across all bioassay observations for pyrethroid insecticides, we obtained a root mean square error (RMSE)^28^ of 0.179 across the out-of-sample predictions of mean proportional mortality for the west and east regions combined (Supplementary Fig. 5). Across all DDT bioassay observations, the corresponding out-of-sample RMSE was 0.167. The individual model constituents of our ensemble included three machine-learning models: an extreme gradient boosting model (XGB), a random forest model (RF) and a boosted generalized additive model (BGAM). We compared the out-of-sample RMSE obtained by the model ensemble to that obtained by each constituent machine-learning model, and confirmed that the prediction error of the Gaussian process meta-model was lower than that of each constituent model (Supplementary Tables 2 & 3). Of the three machine-learning models, XGB had the lowest out-of-sample prediction error followed by RF and then BGAM. The fitted mean model weights given by the Gaussian process metamodel were higher for models with lower out-of-sample prediction error (Supplementary Table 4).

We also performed 10-fold out-of-sample validation to assess the accuracy of the credible intervals of the posterior distributions of predicted mean mortality to pyrethroids. The coverage of the predicted credible intervals was found to be accurate when the measurement error associated with the data, estimated by the Bayesian geostatistical model^29^, was accounted for (Supplementary Fig. 6). Prediction error is heterogeneous across space and time, with the 95% credible intervals of predicted mean mortality being higher in the east compared to the west region (Figs. 4 & Supplementary Fig. 7), and particularly high credible intervals in the north western part of the east region. This reflects the more sparse distribution of bioassay sampling locations in the east region, particularly in South Sudan and much of southern Sudan (see Methods).

### Influential predictor variables

Our models used over 100 potential explanatory variables (see the Methods section), and our results show which of these variables were most influential to the predictions of mean prevalence of resistance. We obtained measures of variable importance for each of the three constituent machine-learning models (XGB, RF and BGAM). Variable importance measures describe the influence of a variable on model predictions relative to the other predictor variables, but they can be hard to interpret when predictor variables are correlated (see Supplementary Note 6), and do not identify causal relationships (see the Methods and Discussion). For each model, the importance of each variable is expressed as a fraction of the total importance across all predictor variables. In ranking variable importance, we weighted the importance of each variable given by each model by the model’s weight obtained from the Gaussian process metamodel for pyrethroids (Supplementary Table 4). This increasingly weights those variables that were more important to models that performed better and thus made a higher relative contribution to the predictions made by the ensemble (Fig. 6). Thus the variable importance values given by XGB and RF are up weighted relative to those given by BGAM. The original variable importance values produced by each model are given in Supplementary Tables 5 and 6 and a description of each predictor variable is given in Supplementary Table 9.

For the west region, variables describing the coverage of insecticide-treated bednets (ITNs) had the highest importance value for each of the three models. For the XGB and RF, the three-year lag of ITN coverage had the highest importance value. For BGAM, non-lagged ITN coverage had the highest importance value and the three-year lag of ITN coverage had the second highest importance value (Fig. 6, Supplementary Table 5 and Supplementary Note 6). Outside the top two, variables describing climate processes, and the area of harvested crops, are highly ranked (within the top 20 most important variables) for all three models (Fig. 6, Supplementary Table 5 and Supplementary Note 6). For the east region, variables describing ITN coverage and rainfall were ranked in the top ten most important variables for all three models (Fig. 6 and Supplementary Table 6). More broadly, variables describing climate processes were highly ranked by all three models. Our ability to quantitatively compare differences in importance across our set of predictor variables is, however, inhibited by differences in the definition of variable importance used in the different machine learning approaches that we have employed (see Methods).

## DISCUSSION

Here, we have quantified spatial and temporal trends in insecticide resistance in the *An. gambiae* species complex in east and west Africa, showing marked increases the prevalence of resistance to pyrethroids and DDT in recent years, as well as geographic expansion. These results highlight the urgency of identifying and implementing effective resistance management strategies. Our predictive maps of mean prevalence of resistance are available to visualise alongside the latest susceptibility test data on the IR mapper website (http://www.irmapper.com), and can guide decisions about resistance management at regional and local levels. In making recommendations, our results will need to be considered in combination with (i) data from resistance monitoring of field samples, including other malaria vector species such as *An. funestus*; (ii) data on the presence of underlying mechanisms of resistance, and (iii) analyses of the expected impacts of resistance management strategies on malaria prevalence^9, 30^. Decision-making frameworks also need to explicitly incorporate predictive uncertainty, which is facilitated by our out-of-sample validation results and our mapped Bayesian credible intervals. Our predictions are not a substitute for ongoing resistance monitoring requirements, but highlight areas with particularly high levels of predictive uncertainty, such as parts of South Sudan, southern Sudan and the Democratic Republic of Congo (Fig. 5D). In these areas, field sampling to measure resistance is the only means of informing resistance management decisions.

**Figure 5.**
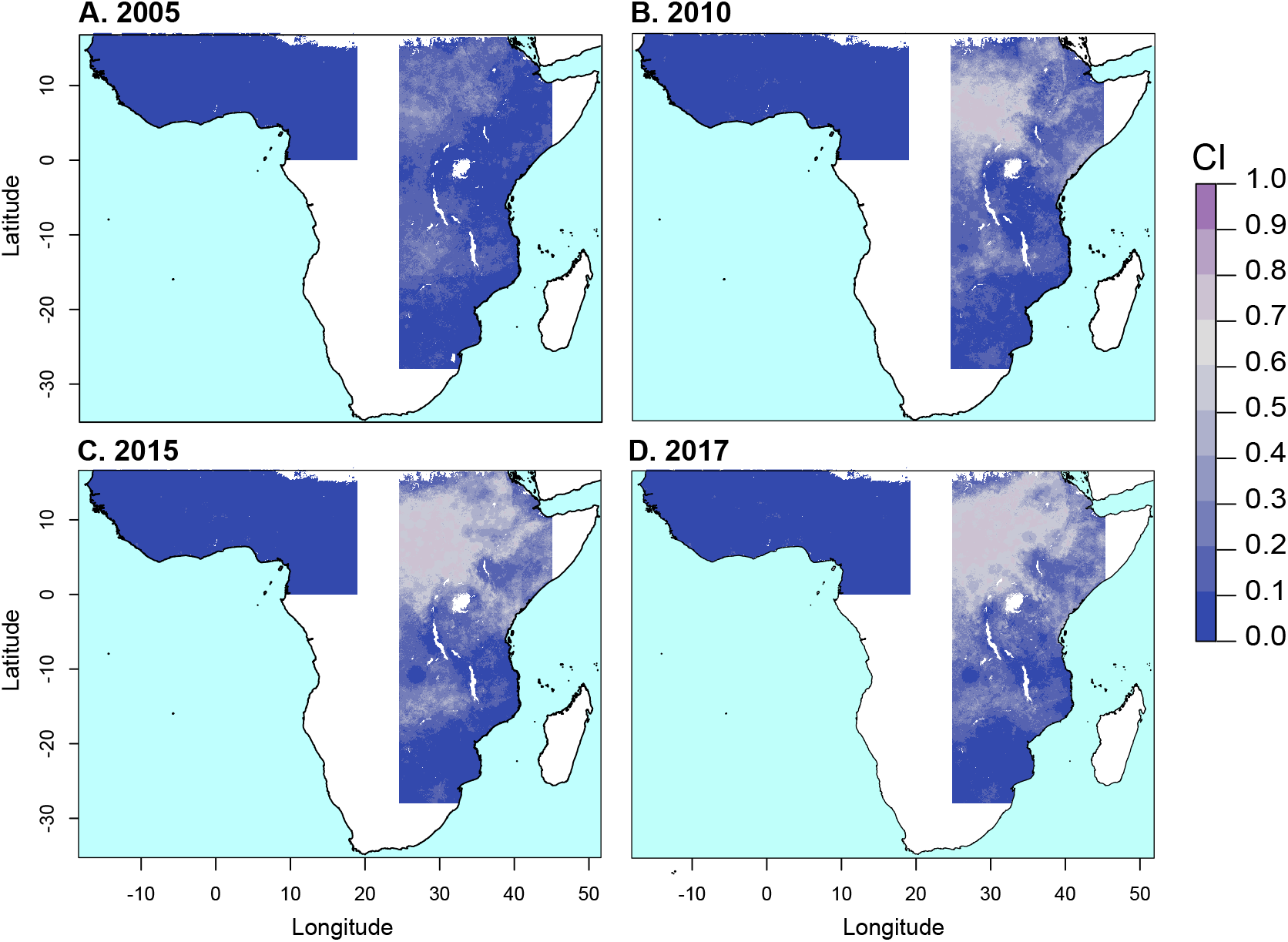
The prediction error (95% credible interval) associated with predicted mean mortality to deltamethrin.

**Figure 6.**
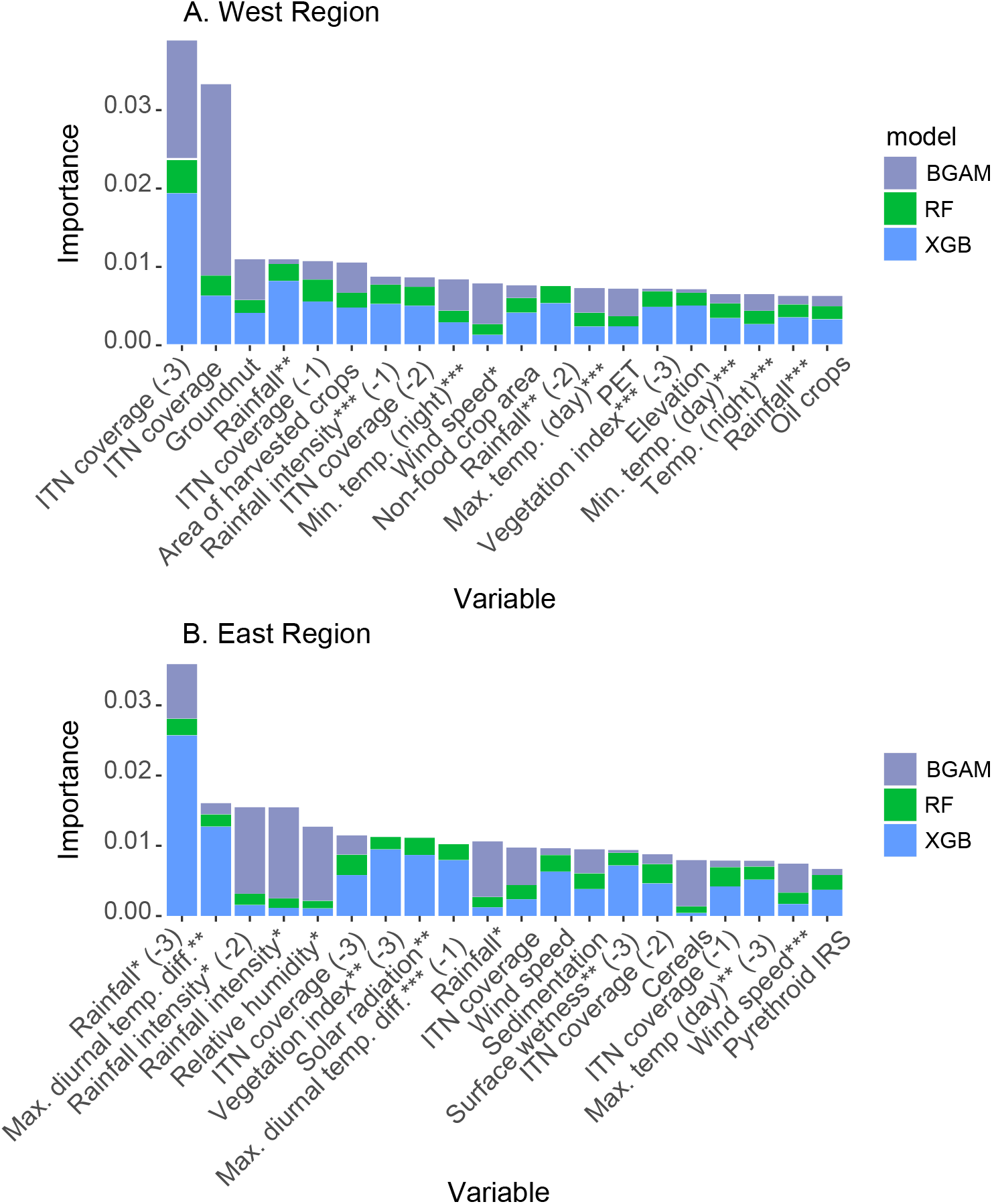
Weighted variable importance of predictor variables given by the three machine-learning models included in the model ensemble for the west (A) and east (B) regions. Stacked bars show the relative variable importance given by the extreme gradient boosting model (blue), the random forest model (green) and the boosted generalized additive model (grey), weighted by the fitted weight for each model given by the Gaussian process meta-model (see text). Variables are ranked by the total height of the stacked bars across the three models, and the top 20 variables are shown. Definitions of each predictor variable are given in Supplementary Table 9. Variable name suffixes (−1), (−2) and (−3) denote time lags of 1, 2 and 3 years, respectively. One, two and three asterisks denote the first, second and third principal component, respectively, for variables available on a monthly time step (see Methods).

Our results show substantial variation in resistance trends between east and west Africa, and within these two regions. Interestingly, ITN coverage was identified as a relatively influential predictor in our models, which is consistent with other studies that have found significant, but spatially variable, increases in pyrethroid resistance associated with the introduction of ITNs^12^. However, in several areas of the central and southern parts of east Africa, such as west Tanzania, ITN coverage has been relatively high (>50%) from 2012-2017^31^ but predicted pyrethroid resistance in 2017 is relatively low (Fig. 1D). This may be influenced by the locations where resistance mechanisms first emerged, patterns of subsequent gene flow including restricted flow across the Rift Valley^32,33,34^, and differences among the sibling species within the *An. gambiae* complex^15, 20^. For example, the distribution of *An. arabiensis* extends further south than other species in the complex^23^ and this species is known to be more plastic in its feeding behaviour, biting outdoors and feeding on cattle^33^. Thus it is possible that selection for resistance in this species lags behind other members of the complex^35, 36, 37^. Our predictions of the prevalence of resistance are based on susceptibility tests that often do not identify the sibling species composition of the *An. gambiae* complex sample that was tested. Our analysis only includes test results that are representative of the original sample collected^8, 27^, and our predictions cannot directly represent variation in the prevalence of resistance due to variation in the composition of sibling species^23,38^. Routine identification of the composition of sibling species in tested samples, and the provision of species-specific mortality values, would improve the capacity of susceptibility test data to inform prediction of resistance.

The coverage of pyrethroid IRS was not amongst the most influential predictors in our models, but only a small fraction of the areas that we modelled (<5% of the west region and <15% of the east region) received pyrethroid IRS between 2005-2017^4^. Thus our results do not imply that IRS is not important in driving the selection of resistance. IRS can, however, be a useful tool to prevent the spread of resistance and mitigate its effects, because the number of options available for IRS mean chemical classes can be rotated through time, applied in a mosaic in space, or combined for use in the same place and time^9^.

It is also important to note that while our models included over 100 potential predictor variables that may influence selection for resistance, it is unlikely that we have captured the full set of causal variables underlying selection. In particular, data on the quantities of insecticides used in agriculture, and where they were applied, was not available^39^. Such information would better inform models of predictive relationships between resistance and agricultural insecticide use. Further, more extensive data on the presence of resistance mechanisms, including a wider coverage of *Vgsc* allele frequencies, as well as metabolic resistance markers^40^, in field populations would potentially aid in predicting and interpreting resistance trends. The similarity in predicted spatiotemporal patterns in resistance across the four pyrethroids and DDT (e.g. Figs. 1 & 3) suggests common underlying resistance mechanisms^27^.

While our analysis focuses on pyrethroids, insecticides from other classes such as carbamates and organophosphates are being increasingly used in IRS interventions^4^. The number of available susceptibility test results for insecticides from these classes is relatively low^8^, and spatiotemporal analyses of resistance would benefit greatly from increasing the frequency and spatial coverage of sampling and testing. Susceptibility test data is also more limited for *An. funestus*, a major malaria vector in Africa that is widespread and among the dominant vector species^41^.

In summary, our results provide an Africa-wide perspective on recent trends in pyrethroid and DDT resistance in *An. gambiae* complex malaria vectors, demonstrating increasingly high prevalence of resistance to the main insecticides used in malaria control. The rapid spread of resistance across large parts of the Sub-Saharan Africa signals an urgent need to quantify the efficacy of different resistance management strategies, and to understand the impact of resistance on malaria transmission and control. Our maps show marked broadscale spatial heterogeneity in resistance, motivating the implementation and assessment of a wide range of strategies that target different insecticide resistance and malaria transmission settings.

## METHODS

### Data

#### Insecticide resistance bioassay data

Insecticide resistance bioassay data were obtained from a published database^8^, which is an updated version of the data used in Hancock et al.^27^ that includes samples tested up until the end of 2017. The data record the number of mosquitoes in the sample and the proportional sample mortality resulting from the bioassay, as well as variables describing the mosquitoes tested, the sample collection site, and the bioassay conditions and protocol. We used this information to select a subset of records for inclusion in our study (Supplementary Note 7). In summary, we include bioassay results where standard WHO susceptibility tests or CDC bottle bioassays using either one of the four pyrethroid types (deltamethrin, permethrin, lambda-cyhalothrin and alpha-cypermethrin) or the organochlorine DDT were performed on mosquito samples belonging to the *An. gambiae* species complex. We include results from bioassays conducted over the period 2005-2017. Due to spatial heterogeneity in the sampling distribution we confine our analysis to samples collected from within two separate geographic (west and east) regions of Sub-Saharan Africa (see Supplementray Fig. 11 and Supplementary Note 7). We excluded Madagascar from our analysis, as our models of resistance on the mainland may not generalize well to island populations. The final number of proportional mortality observations across all insecticide types was 6423 across 1466 locations, with 3515 and 2908 observations in the west region and east region, respectively (Supplementary Tables 7 & 8).

#### Voltage-gated sodium channel (Vgsc) allele frequency data

The *Vgsc* is the target site for both pyrethroids and DDT and mutations in this channel confer resistance. Our analysis used data on the frequency of *Vgsc* mutations in mosquito samples belonging to the *An. gambiae* species complex collected from within the west and east regions over the period 2005-2017^8, 27^. These data record the combined frequency of the single point mutations L1014F and L1014S with respect to the wild type allele L1014L, and comprise 316 observations (215 observations for the west region and 101 observations for the east regions; Supplementary Table 7). As described below, we incorporated these data into machine learning models in order to inform prediction of phenotypic resistance to DDT and pyrethroids by exploiting the positive association between the frequency of *Vgsc* mutations and the prevalence of these resistance phenotypes^27^.

#### Potential predictor variables

Our set of predictors includes 111 variables describing environmental characteristics that could potentially be related to the development and spread of insecticide resistance in populations of Gambiae complex mosquito species (described in Supplementary Table 9 and Supplementary Note 8). These variables describe the coverage of insecticide-based vector control interventions, agricultural land use^42 43^ and the environmental fate of agricultural insecticides^39^, other types of land use^42, 44, 45, 46^, climate^42, 47, 48^, and relative species abundance. Our vector control intervention data includes a variable estimating the yearly coverage of insecticide-treated bed nets (ITNs)^31,49^ and a variable estimating the coverage of indoor residual spraying (IRS) with either pyrethroids or DDT year^4^. Relative species abundance is represented by a variable estimating the abundance of *An. arabiensis* relative to the abundance of *An. gambiae* and *An. coluzzii*^38^. For all variables, we obtained spatially explicit data on a grid with a 2.5 arc-minute resolution (which is approximately 5 km at the equator) covering Sub-Saharan Africa. For variables for which temporal data were available on an annual resolution, we included time-lagged representations with lags of 0, 1, 2 and 3 years.

### Gaussian process stacked generalization ensemble modelling approach

Stacked generalization is a method of combining an ensemble of models to produce a meta-model, with the aim of achieving better predictive performance than the individual model constituents^50, 51^. Here we adopt a stacking design whereby a set of individual models that make up the first layer, referred to as the *level 0 models*, feed into a single meta-model on the second layer, referred to as the *level 1 model*. We use the Gaussian process stacked generalization approach developed by Bhatt et al.^25^, which uses Gaussian process regression as the level 1 model that combines weighted out-of-sample predictions from a set of multiple level 0 models derived from machine learning methods. The approach exploits the known strengths of these different methodologies, using machine learning methods to extract as much predictive power from the covariates as possible, and then allowing the Gaussian process to model the spatiotemporal error covariance structure, aiming to further improve prediction. Bhatt et al.^25^ showed that, under the (restrictive) assumption that the true function is a part of the models function space, the use of the Gaussian process model of residual variation improves prediction accuracy compared to a standard constrained weighted mean across the ensemble predictions.

#### Machine learning models

Our set of level 0 models consists of three different types of machine learning model that predict insecticide resistance, using our bioassay mortality observations as the label and our suite of intervention, agriculture and environmental covariates as features. The machine learning approaches employed include extreme gradient boosting (implemented using the R package xgboost), random forests (implemented using the R package randomForest), and boosted generalized additive models (implemented using the R package mboost). We chose these methods because of their demonstrated high predictive performance, particularly in previous applications of Gaussian process stacked generalization to spatial processes^25^. The label for the level 0 models was the proportional mortality observations from bioassays conducted using the four pyrethroid types (deltamethrin, permethrin, lambda-cyhalothrin and alpha-cypermethrin), the proportional mortality observations for bioassays conducted using DDT, and the observations of the combined frequency of the *Vgsc* mutations L1014F and L1014S. We included in the label our data on the observed combined frequency of *Vgsc* mutations in mosquito samples, because these observations are significantly associated with the prevalence of resistance to DDT and pyrethroids^27^, and can therefore inform prediction of these mortality values. Before performing parameter tuning on the level 0 models we applied two data transformations to the label, the empirical logit transformation followed by the inverse hyperbolic sine (IHS) transformation^52^.

The features used in the models included the 111 environmental predictor variables together with the one, two and three year lags for those variables that vary temporally (on a yearly time step). A factor variable grouping the label according to the type of observation was also included as a feature, assigning a different group to bioassay observations depending on type of insecticide used and whether a WHO or CDC susceptibility test was used. This factor variable also assigned the *Vgsc* allele frequency observations to a separate group. Finally, the year in which the bioassay and allele frequency samples were collected was also included as a feature.

For each level 0 model, parameter tuning was performed using *K*-fold out-of-sample validation based on subdividing the data into *K* training and validation subsets (see Supplementary Note 7). In applying the extreme gradient boosting method we used the DART boosting methodology to avoid overfitting^53^.

#### Model stacking and Gaussian process regression

Let *g_A_*(**s**_*i*_, *t*) denote the (empirical logit and IHS transformed) proportional mortality record for a bioassay using insecticide type *A* conducted on a sample collected at geographic coordinates **s**_*i*_ and sampling time *t*. To implement Gaussian process stacked generalization, we model the transformed observations, denoted *g_A_*(**s**_*i*_,*t*), using a Gaussian process regression formulation:

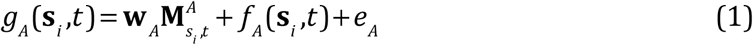

where **w**_*A*_ is a constant vector, 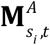 is a design matrix, *f_A_*(**s**,*t*) is a Gaussian process modelled by a spatiotemporal Gaussian Markov random field (GMRF)^54^, and *e_A_* is Gaussian white noise 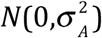. We define a Bayesian hierarchical formulation for the model (eqn 1) using a vector of prior probability distributions for the hyperparameters *θ_A_* = [**w***_A_*, *ψ_A_*, *σ_A_*] where *ψ_A_* are the parameters of *f_A_*(**s**,*t*) (see Supplementary Note 6). To fit the model, the elements of the design matrix 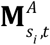 are set to the out-of-sample predictions of the level 0 models derived from *K*-fold cross-validation i.e. 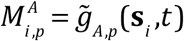, where 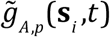 is the prediction of the *i^ith^* withheld (transformed) observation *g_A_*(**s**_*i*_.,*t*) given by the *p^th^* level 0 model. Validation folds were randomly selected from the full data set. Posterior distributions of *θ_A_* and *f_A_*(**s**,t) are then estimated by fitting the model (eqn 1) using the R-INLA package (www.r-inla.org)^55^. The posterior mean of the vector **w**_*A*_ contains the fitted weights for each model, representing the relative contribution of each model to the predictions made by the model ensemble. Our implementation of Gaussian process regression (eqn 1) constrains each weight to be positive (*w_p_* ≥ 0,∀*p*)^56^. Once the parameter estimation has been performed, the final set of predictions, 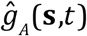, given by the stacked model are obtained by replacing the elements of 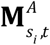 with the in-sample predictions of the 10 models obtained by fitting each of these models to the all the data (all the labels and the corresponding sets of features)^25^ (Supplementary Notes 6 & 7).

#### Posterior validation

We performed posterior validation of the stacked model using 10-fold out of sample cross-validation (withholding each validation fold from both the level 0 and level 1 models). We used these out-of-sample predictions to assess the accuracy of the predicted means of the observations as well as their predicted credible intervals (Supplementary Note 7). We also assessed the suitability of our assumed data generating process using probability integral transform (PIT) histograms on out-of-sample data (Supplementary Note 3).

#### Predictor variable importance

We calculated measures of the importance of each predictor variable for each of the machine learning models used in our model ensemble. For the extreme gradient boosting model we used the gain measure calculated for each variable using the xgboost package^57^, which is the fractional total reduction in the training error gained across all of that variable’s splits. For the random forest model we use the permutation importance measure calculated using the randomForest package^58^, which is the fractional change in the out-of-bag error when the variable is randomly permuted. In the case of the boosted generalized additive model, we use the mboost package^59^ to calculate variable importance as the total reduction in the training error across all boosting iterations where that variable was chosen as the base learner. For each model, we express the importance of a single variable as a fraction of the total importance across all predictor variables in that model.

## DATA AVAILABILITY

The predictive maps of the mean prevalence of resistance are available to download from Figshare (https://figshare.com/s/00b829f256694ed3c632) and will be available to visualise on the Malaria Atlas Project website (https://map.ox.ac.uk/explorer/#). The susceptibility test data is available to download (https://doi.org/10.1101/582510^8^). Sets of susceptibility test data and predictor variable data in the form used by the statistical modelling analyses are available from GitHub.

## CODE AVAILABILITY

R code for implementing the extreme gradient boosting, random forest, and boosted generalized additive models and the R-INLA geostatistical models is available on GitHub.

## ACKNOWLEDGEMENTS

The authors are extremely grateful to the many people who contributed unpublished datasets and to the authors who provided additional information linked to their published works. This work was funded by Wellcome Trust Grant 108440/Z/15/Z (to C.L.M.).

## AUTHOR CONTRIBUTIONS

P.A.H., C.J.M.H., M.C., P.W.G and C.L.M. designed the analyses; P.A.H. and C.L.M. led the writing of the manuscript; P.A.H. performed the statistical modelling analyses; C.J.M.H., J.T., H.G., S.B. and C.L.M. contributed data layers for the predictor variables used in the statistical models; E.C., S.B., P.W.G. and C.L.M. advised the statistical modelling analyses; C.J.M.H., H.G., J.H., M.C., and S.B. contributed to writing the manuscript.

